# Visual evidence accumulation guides decision-making in unrestrained mice

**DOI:** 10.1101/195792

**Authors:** Onyekachi Odoemene, Sashank Pisupati, Hien Nguyen, Anne K. Churchland

**Affiliations:** Watson School of Biological Sciences; Cold Spring Harbor Laboratory, Cold Spring Harbor NY 11724

## Abstract

The ability to manipulate neural activity with precision is an asset in uncovering neural circuits for decision-making. Diverse tools for manipulating neurons are available for mice, but the feasibility of mice for decision-making studies remains unclear, especially when decisions require accumulating visual evidence. For example, whether mice’ decisions reflect leaky accumulation is not established, and the relevant and irrelevant factors that influence decisions are unknown. Further, causal circuits for visual evidence accumulation have not been established. To address these issues, we measured >500,000 decisions in 27 mice trained to judge the fluctuating rate of a sequence of flashes. Information throughout the 1000ms trial influenced choice, but early information was most influential. This suggests that information persists in neural circuits for ~1000ms with minimal accumulation leak. Further, while animals primarily based decisions on current stimulus rate, they were unable to entirely suppress additional factors: total stimulus brightness and the previous trial’s outcome. Next, we optogenetically inhibited anteromedial (AM) visual area using JAWS. Importantly, light activation biased choices in both injected and uninjected animals, demonstrating that light alone influences behavior. By varying stimulus-response contingency while holding stimulated hemisphere constant, we surmounted this obstacle to demonstrate that AM suppression biases decisions. By leveraging a large dataset to quantitatively characterize decision-making behavior, we establish mice as suitable for neural circuit manipulation studies, including the one here. Further, by demonstrating that mice accumulate visual evidence, we demonstrate that this strategy for reducing uncertainty in decision-making is employed by animals with diverse visual systems.

**Significance statement:** To connect behaviors to their underlying neural mechanism, a deep understanding of the behavioral strategy is needed. This understanding is incomplete in mouse studies, in part because existing datasets have been too small to quantitatively characterize decision-making behavior. To surmount this, we measured the outcome of over 500,000 decisions made by 27 mice trained to judge visual stimuli. Our analyses offer new insights into mice’ decision-making strategies and compares them with those of other species. We then disrupted neural activity in a candidate neural structure and examined the effect on decisions. Our findings establish mice as a suitable organism for visual accumulation of evidence decisions. Further, the results highlight similarities in decision-making strategies across very different species.

## Introduction

Rodents have emerged as a powerful model organism for probing the neural circuits underlying decision-making (Carandini and Churchland 2013). Mice are an ideal model for studying neural circuits because of the tools for accessing and probing genetically defined cell types (Taniguchi et al. 2011; Madisen et al. 2010, 2012, 2015). Despite these advantages, other species are more commonly used in perceptual decision-making studies that involve temporal accumulation of sensory evidence, perhaps due to the assumption that such tasks are too difficult for mice. However, mice have been trained on numerous sensory perception tasks (Andermann et al. 2010; Busse et al. 2011; Sanders and Kepecs 2012; Glickfeld et al. 2013; Guo et al. 2014a; Burgess et al. 2017; Goard et al. 2016; Marbach and Zador 2016; Funamizu et al. 2016; Jeong et al. 2017). This suggests that they might be suitable for visual evidence accumulation tasks, and several studies report promising performance in mice on such tasks (Douglas et al. 2006; Stirman et al. 2016; Morcos and Harvey 2016).

Two major gaps in our understanding of accumulation of evidence decisions are apparent: a precise characterization of the timecourse of accumulation, and an understanding of how relevant and irrelevant factors jointly shape decisions. Recent work suggested that a sequence of visual cues collectively influence an eventual decision (Morcos and Harvey 2016). This data constitutes an essential first step in establishing mice as a suitable decision-making model, but outstanding questions remain. First, a large-scale study with many subjects and large trial counts is needed. This benefits a number of analyses, including those that evaluate the influence on the final decision of stimuli arriving at different times. Such analyses require many trials to achieve the level of precision that is required to fully characterize the evidence accumulation timecourse. Further, many animals are required to distinguish idiosyncratic strategies from the overall tendency of the species. For instance, in existing work (Morcos & Harvey 2016), the use of 5 mice hinted that the most common strategy is to weight early evidence over late evidence, but the small animal number and the variability in strategy made a firm conclusion difficult. Closing these gaps in our understanding of accumulation decisions is essential, especially for interpreting causal manipulations (Krakauer et al. 2017).

Indeed, the causal circuits for visual evidence accumulation are not established in mice although inactivations have demonstrated a role for cortical and subcortical structures in other kinds of mouse decisions. For instance, suppressing the posterior parietal cortex (PPC) in mice impairs memory-guided decisions (Harvey et al. 2012; Funamizu et al. 2016; Goard et al. 2016). These studies suggest a role for cortical circuits in decision-making and visually guided behavior. However, the role of these structures in visual evidence accumulation is unknown. The importance of establishing a causal role for a putative neural structure is underscored by recent results that even areas strongly modulated during behavior might not be part of the causal circuit (Katz et al. 2016; Erlich et al. 2015).

Overall, mice have potential as an animal model for decision-making, but have been held back because of lack of detailed knowledge about behavior and little insight into the contribution of cortical circuitry. Here, we begin to close those gaps. First we report that mice accumulate evidence for visual decisions. Our large dataset and use of stochastic stimuli allow precise characterize how information presented at different times influences decisions. Next, we report that mice’ decisions are jointly shaped by stimulus rate, brightness and the outcome of previous trials. Two independent experiments demonstrated that the influence of brightness was larger in rats compared to mice. Finally, we report that suppressing activity in the Anteromedial visual area (AM) biases decisions, highlighting the role of cortex in evidence accumulation. Taken together, these findings establish mice as a suitable organism for visual accumulation of evidence decisions. Further, the results highlight similarities in decision-making strategies across very different species.

## Materials and Methods

### Animal Subjects

The Cold Spring Harbor Laboratory Animal Care and Use Committee approved all animal procedures and experiments and all experimental procedures were in accordance with the National Institutes of Health’s *Guide for the Care and Use of Laboratory Animals*. Experiments were conducted with female or male mice between the ages of 6-25 weeks. All mouse strains were of C57BL/6J background and purchased from Jackson Laboratory. Ten GCaMP6f transgenic mice (Ai93 /Emx1-cre /CamKIIα-tTA) of both sexes were used for retinotopic mapping and area AM photoinhibition experiments. Four male Long Evan rats (6 weeks, Taconic) were also used for behavior experiments.

### Behavioral Training

Before behavioral training, mice were gradually water restricted over the course of a week. Mice were weighed daily and checked for signs of dehydration throughout training period (Guo et al. 2014b). Mice that weighed less than 80% of their original pre-training weight were supplemented with additional water. Behavioral training sessions lasted 1-2 hours during which mice typically harvested at least 1 mL of water. Mice rested on the weekends. Mice who failed to harvest at least 0.4 mL on two consecutive days were supplemented with additional water.

Animal training took place in a sound isolation chamber containing a three port setup described elsewhere (Raposo et al. 2012). Mice poked their snouts into the center port to initiate trials and trigger sensory stimuli. Animals reported choices by moving to a left- or right-side port. In the first training stage, mice learned to wait for at least 1100 ms at the center port before reporting their decision. We shaped the behavior by rewarding the mice at the center port (0.5 μL) and gradually increasing the minimum wait duration from 25 ms to 1100 ms over the course of 1-2 behavioral sessions. Without center reward, this stage typically took 10-12 sessions to learn.

During the first stage, mice were not rewarded for making the correct association between the stimulus and response port; rather, on each trial, a random port (left or right) was chosen as the reward port and a liquid reward (2 to 4μL) was delivered to the port. Trials in which the mouse waited the minimum required duration at the center port are referred to as completed trials.

In the second stage of training, mice learned to associate high-rate flash sequences (13-20 flashes/s) with the right port and low-rate flashes (1-11 flashes/s) with the left port. Trials with 12 flashes/s were randomly rewarded. For some mice, the contingency was reversed, such that high-rate flashes were rewarded at the left-hand port and low-rate flashes were rewarded at the right-hand port. Mice received a liquid reward for correct responses. They were punished for incorrect responses or for withdrawing early with a time-out period (2 to 4 s), during which they could not initiate a trial.

We employed several anti-bias methods to correct the side bias, which often occurred when mice began stage two. Anti-bias strategies included: physically obstructing access to the biased port, changing the reward size, and modifying the proportion of left vs. right trials.

Training was considered complete when mice waited at least 1100ms at the center port, performed above 80% percent correct on the easiest flash rates (Figure 1B, C), and experienced at least 8 or more flash rates. This required approximately 2 months, with one daily session 5 days per week.

**Figure 1.**
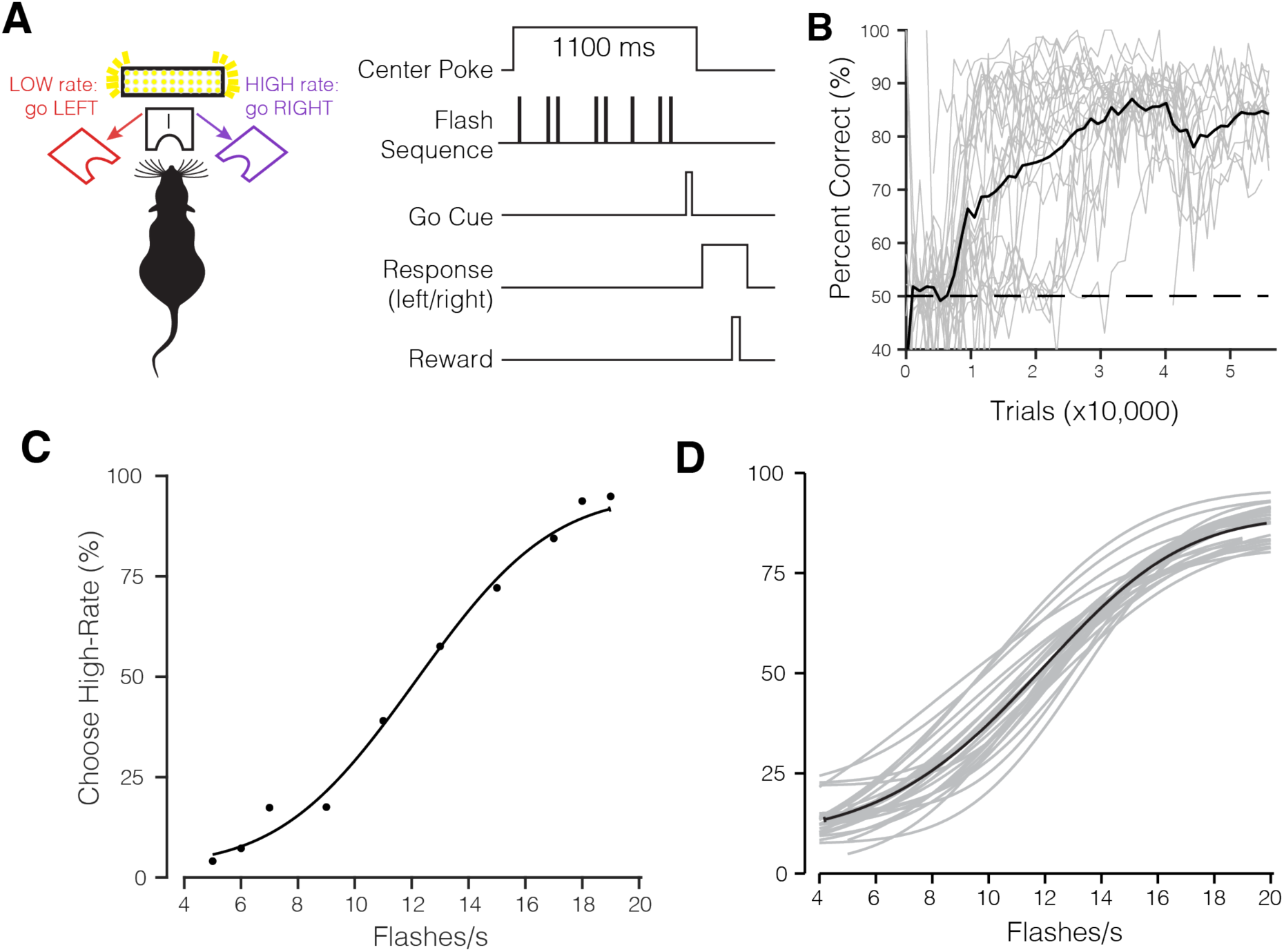
A dataset of over half a million trials demonstrates mice can be trained to make stable and reliable decisions about visual stimuli. (A) Task schematic and trial structure of the three-port choice task. The mouse initiated trials and stimulus delivery by poking the center port. Mice reported whether stimuli were low-rate (left port) or high-rate (right port). Mice waited at the center port for at least 1100 ms, with the stimulus delivered after a variable delay (10-100ms). At the end of the 1000ms stimulus period, an auditory “Go” tone was played. Correct choices to the left or right were rewarded with a small drop of water (2 μL), incorrect choices were followed by a 2-3 s timeout. (B) Percent correct on easiest stimulus conditions (4 and 20 flashes/s) plotted across total trials experienced by the mouse. Individual mice: gray traces and average: black trace, 27 mice. (C) Psychometric function fit for individual mouse from single session (494 trials), and (D) Pooled data from 27 mice averaged across multiple sessions (537,288 trials). Individual mice: gray traces and average: black trace.

Stimulus presentation, reward delivery, and data collection were performed through a MATLAB interface and Arduino-powered device (BPod, SanWorks LLC).

### Visual stimuli

Stimuli were sequences of 20 ms pulses of light from a LED panel (Ala Scientific). The inter-pulse intervals were randomly selected from a discrete exponential distribution (Brunton et al. 2013). For the exponential interval stimulus, the minimum inter-pulse interval was 20 ms, and the number of flashes for a given stimulus determined the maximum interval. 4-20 flashes/s were presented on each trial, always over the course of 1000ms. The stimulus was created using 25 fixed time bins each 20ms in duration. A Poisson coin flip determined whether an event (flash) would occur in each bin. An empty 20ms time bin followed each flash. This 1000ms period was followed by a 100 ms delay, leading to a total time in the port of 1100ms.

Each 20ms flash pulse was generated by a half-wave rectified sinusoidal signal thresholded at the peaks and with a base frequency of 200Hz. This approach effectively controls the total LED on-time or the “density” of the 20ms pulses. It is similar to pulse-width modulation technique used to control LED brightness. During normal sessions, the base frequency is multiplied by a brightness factor, which is kept constant across sessions.

### Brightness Manipulation

For the brightness manipulation experiments, the 20ms flash duration was held constant, so that the subjects could not use the flash duration as a cue for the correct stimulus category. In the uniform brightness manipulation experiment, the normal brightness factor was either halved or doubled to produce the “dimmer” and “brighter” conditions. In the uncorrelated brightness manipulation experiment, we varied the LED on-time within the flash duration such that the brightness factor was inversely scaled with the flash rate. Because the lowest number of flashes presented was 4 flashes/s, and we did not change the flash duration, we normalized all flash sequences such that the total LED on-time was equal to 4 flashes/s. All brightness manipulations were randomly introduced on 5% of all trials.

### Head bar implantation and skull preparation

For retinotopic mapping experiments, mice were implanted with a custom titanium head bar. Mice were anesthetized with isoflurane (2%) mixed with oxygen and secured onto a stereotaxic apparatus. Body temperature was maintained at 37°C with a rectal temperature probe. The eyes were lubricated with Puralube ointment before the start of the surgery, followed by subcutaneous injection of analgesia (Meloxicam, 2mg/kg) and antibiotic (Enrofloxacin, 2mg/kg). Fur on the scalp was removed with hair clippers and Nair (Sensitive Formula with Green Tea), followed by betadine (5%) swab. Lidocaine (100 μL) was injected underneath the scalp before removing the scalp. The skull was cleaned with saline and allowed to dry. A generous amount of Vetbond tissue glue (3M) was then applied to seal the skull. Once the Vetbond was dry, the head bar was secured with Metabond (Parkwell) and dental acrylic. Postsurgical analgesia was applied. Mice were allowed 3 days to recover before retinotopic mapping.

### Retinotopic Mapping

Retinotopic mapping was performed in awake head-fixed animals adapted from (Garrett et al, 2014; Juavinett et al. 2016). Periodic (Fourier) stimulation was used: a narrow bar (10°) was drifted across the four cardinal directions of the screen. Presented within the drifting bar was a flickering checkerboard pattern (12° checks, 5Hz). One trial consisted of 11 sweeps of the bar in 22 seconds in one of the four cardinal directions. The first cycle was discarded because it introduced stimulus onset transients. Each trial was repeated 15 times for each direction. The monitor was placed in the visual hemifield contralateral to the imaging hemisphere, positioned at an angle of 77° from the midline of the mouse and a distance of 15 cm. Imaging data was acquired at 20 frames per second.

### Optogenetic Inactivation

For JAWS inhibition experiments, mice were injected with AAV8-CamKII-JAWS-KGC-GFP-ER2 (UNC Vector Core) into area AM identified by retinotopic mapping. AM is a prominent candidate for decision-making as it appears to at least partially overlap with the previously defined location of posterior parietal cortex (Funamizu et al. 2016; Krumin et al. 2017) and has projections to frontal and motor areas. Similar projection patterns have been observed in primate lateral intraparietal area LIP (Cavada and Goldman-Rakic 1989a, 1989b), an area routinely implicated in perceptual decision-making studies (Gold and Shadlen 2007; Hanks and Summerfield 2017).

Virus injections were performed using Drummond Nanoject III, which enables automated delivery of small volumes of virus. To minimize virus spread, the Nanoject was programmed to inject slowly: six 30 nL boluses, 60 s apart, and each bolus delivered at 10 nL/sec. Approximately 180nL of virus was injected at multiple depths (200 and 500 μm) below the brain surface. Following the virus injection, 200 μm fiber (metal ferrule, ThorLabs) was implanted above the injection site. The optical fiber was secured onto the skull with Vitrebond, Metabond, and dental acrylic. The animals were allowed at least 3 days to recover before behavioral training. A red 640nm fiber-coupled laser (OptoEngine) was used for inactivation. Experiments were conducted with multiple laser power levels: 0.5, 1, and 2 mW (16, 32, and 64 mW/mm^2^). One power level was used per session. On inactivation sessions, laser light was externally triggered using a PulsePal (Sanworks LLC) device. The laser stimulation pattern was a square pulse (1 second long) followed by a linear ramp (0.25s), which began at the onset of the stimulus. Stimulation occurred on 25% of trials.

### Psychometric function

We fitted a four-parameter psychometric function to the responses of subjects that performed the visual flashes categorization task. The general form of the psychometric function defines the probability (*p_H_*) that the subject chooses the port associated with high flash rate as:

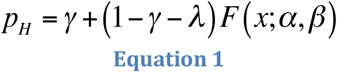

where *γ* and *λ* are the lower and upper asymptote of the psychometric function, which parameterize the guess rate and lapse rate, respectively; *F* is a sigmoidal function, in our case a cumulative Normal distribution; *x* is the event rate i.e. the number of flashes presented during the one second stimulus period; *α* parameterizes the horizontal shift or bias of the psychometric function and *β* describes the slope or sensitivity. The psychometric function *F*(*x; α,β*) for a cumulative Normal distribution is defined as:

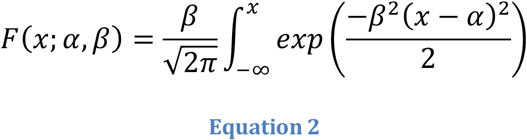

The parameters of the psychometric function were estimated with the Palamedes Toolbox (Prins and Kingdom 2009).

### Choice History

We implemented two probabilistic choice history models to evaluate the influence of prior choice(s) on the current choice of the subject. The first approach, assessed whether success or failure on the most recent trial influenced the performance on the current trial (Busse et al. 2011):

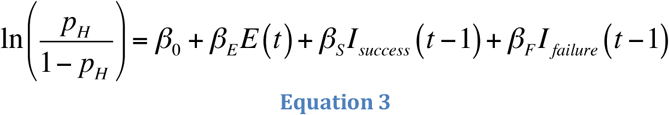

where *t* indicates the current trial and *E* is the signed stimulus evidence of the current trial. Evidence is computed as the difference between the flash rate of the trial and the category boundary (12 flashes/s). *I_sucess_* and *I_failure_* are indicator variables for success (reward) and failure on the previous trial, respectively. The coefficients (*β_0_*, *β_E_ β_S_, and β_F_* were estimated with MATLAB *glmfit*.

The second model (Fründ et al. 2014) assessed the influence of choices made several trials in the past:

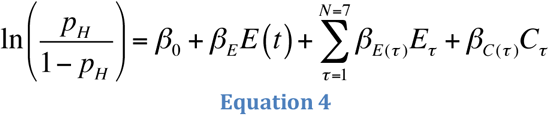

where *t* indicates the current trial and *E*is the signed stimulus evidence of that trial. The additional regressors *E_τ_* and *C_τ_* represent the evidence and choice on *τ* previous trials in the past, respectively. Coefficients (*β_0_*, *β_E_,β_E(τ)_and β_c(τ)_*) were estimated with MATLAB *glmfit*.

### Logistic Regression Reverse Correlation

The logistic regression function for estimating the weights associated with each moment of the stimulus (the psychophysical kernel) can be written as:

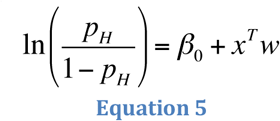

where *β_0_* is a scalar bias term, *x* is a vector of the 25 successive time windows over the trial, and *w* is a vector of the weights for each time window. *β_0_* and weight vector *w* were estimated with the MATLAB function *glmfit*.

### Generalized Linear Mixed Model (GLMM)

To statistically test whether there was a significant effect of photoinhibition of area AM on the population group level, we used a Generalized Linear Mixed-Model (GLMM). GLMMs are an extension of the Generalized Linear Model, which can be used to model both fixed and random effects in categorical data. In psychophysics, GLMMs can be used to generalize results across multiple subjects and experimental conditions (Knoblach and Maloney 2012; Moscatelli et al. 2012; Erlich et al. 2015).

The GLMM model written in the Wilkinson notation:

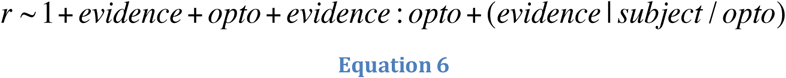

Each term of the equation has a coefficient, *β*. The model specifies that the subject’s response, *r*, is a function of the fixed effects: the *evidence*, defined as the difference between flash rate and the category boundary; its coefficient represents the slope of the psychometric function, the photoinhibition indicator variable *opto*, and the interaction between the *evidence* and *opto*. The interaction term *evidence:opto* evaluates whether photoinhibition alters the subject’s sensitivity or the slope of the psychometric function. The model allows the four fixed effects parameters to vary for each individual subject (random effects). The model uses a probit linking function and was fit using a Maximum Likelihood procedure. The GLMM analysis was performed using the R package *’lme4’* as described in Erlich et al (2015).

The effect of photoinhibition on the horizontal location of the psychometric function was quantified by the choice bias. The choice bias was defined as:

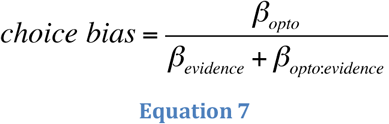

where *β_opto_, β_evidence_, β_evidence:opto_* are estimated coefficients from the GLMM equation above. The choice bias reflects the equivalent change in the stimulus that would recapitulate the observed effects of photoinhibition and is in units of flashes/s. Positive choice bias would indicate that on photoinhibition trials caused the subject to be biased towards high-rate responses. Since the choice bias is computed from estimated parameters of the GLMM model, we computed the errors (95% confidence intervals) via error propagation.

## Results

We trained mice to categorize a stochastic pulsatile sequence of visual flashes (Figure 1), similar to earlier studies with rats (Raposo et al. 2012; Brunton et al. 2013; Scott et al. 2015). Mice performed a three-port choice task (Uchida and Mainen 2003), in which they judged whether the total number of full-field flashes presented during a 1000ms period exceeded an experimenter-defined category boundary (12 flashes/s, Figure 1A). Each flash in the sequence was 20 ms long and followed by an inter-flash interval drawn from a discrete exponential distribution.

### Mice learned to categorize stochastic sequences of visual flashes

Mice performed hundreds of trials per session (median 767 trials) and reached high performance accuracy at the easiest level of the task (Figure 1B). Behavioral performance was quantified by fitting a psychometric function (Eq. 1, Figure 1C,D). Individual mice on single sessions (Figure 1C) and across multiple sessions (Figure 1D) made increasingly more high-rate choices as the flash rate increased, achieving psychometric performance comparable to rats trained on the same task (Raposo et al. 2012).

### Mice decisions were influenced most by flashes early in the sequence

To maximize accuracy, animals should count all the flashes presented during the fixed stimulus presentation period. Because all flashes in the sequence are equally informative about the overall count, subjects should apply an equal weight to all flashes. However, mice might instead make use of only a portion of the stimulus. Overweighting flashes early in the sequence would reflect an impulsive strategy of making up one’s mind too early, whereas overweighting flashes later in the sequence would reflect a forgetful/leaky strategy (Kiani et al. 2008).

To distinguish these strategies, we used the well-established logistic regression approach to estimate the psychophysical kernel (Huk and Shadlen 2005; Katz et al. 2016; Yates et al. 2017). The logistic regression-based reverse correlation approach reveals the timecourse of how incoming stimuli, on average, influence the subject’s choice. Our use of stimuli that appear stochastically over time and our large dataset together enabled a continuous and precise characterization of this timecourse. Across mice (Figure 2A), the entire sequence of flashes was informative, as indicated by non-zero regression weights throughout the trial. Interestingly, flashes presented earlier in the sequence informed the choice more strongly than flashes presented later in the sequence. This implies that mice tended to overweight stimuli presented early in the trial, consistent with an impulsive integration strategy. The psychophysical kernels of rats (Figure 2B) trained on the same task were generally flat, reflecting an integration strategy in which evidence is weighted equally over time.

**Figure 2.**
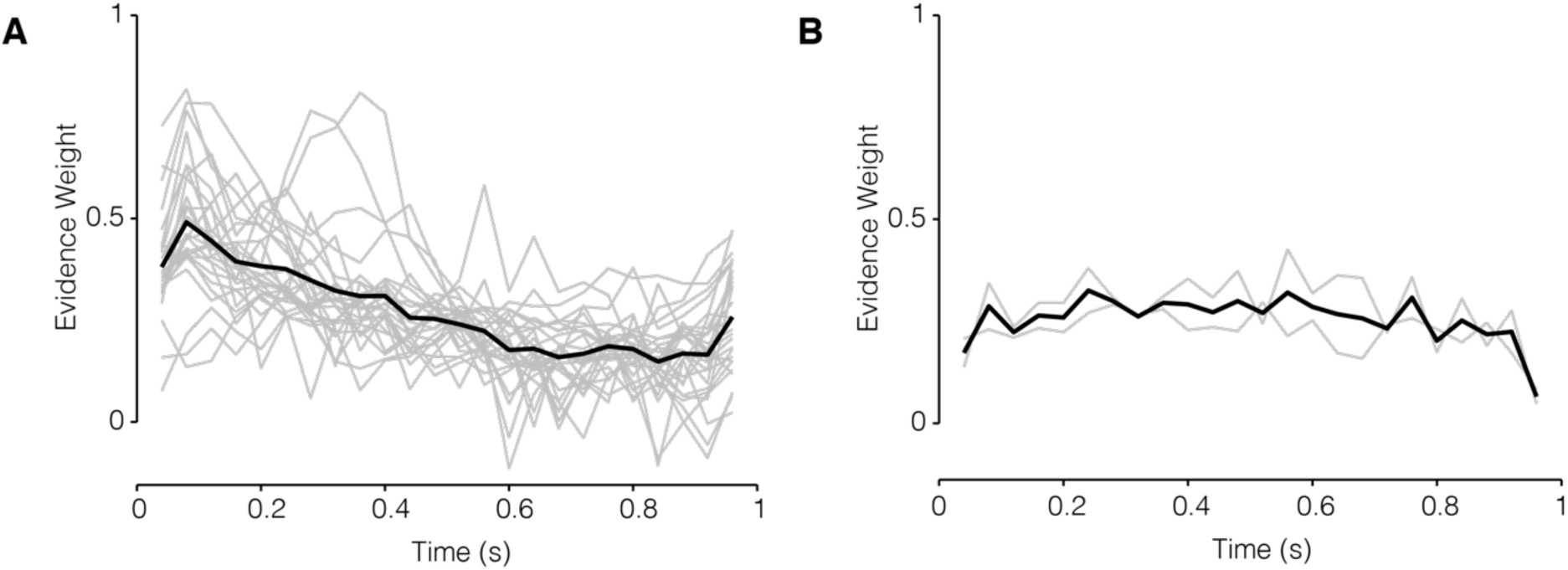
Decisions in mice and rats reflect evidence presented throughout the trial. (A) Psychophysical kernels from 27 mice (537,288 trials) and (B) 2 rats (26,890 trials). Gray traces, individual subjects; black trace, average. Values were above 0 throughout the trial for almost all subjects, demonstrating that stimuli presented throughout the 1000ms duration influence the animal’s eventual choice.

### Mice were influenced by performance on previous trial

Next we used the same dataset to evaluate whether mice were influenced by performance on previous trials. Several studies have reported that human and animal subjects performing perceptual tasks are influenced by previous choices (Busse et al. 2011; Fründ et al. 2014; Scott et al. 2015; Abrahamyan et al. 2016; Urai et al. 2017, Hwang et al. 2017), even when the trials are independent. We used two quantitative models to assess whether the event-based, visual accumulation decisions used here were likewise influenced by choices made on previous trials.

The first approach assessed whether success or failure on the most recent trial influenced the performance on the current trial (Methods; Busse et al. 2011). Figure 3A shows a scatter plot of the coefficients for previous success (β_S_) and previous failures (β_F_). Nearly all the 27 mice had positive β_S_ coefficients, indicating that mice tended to repeat the same choice on the current trial if they were rewarded on the previous trial. Many of the mice also had positive β_F_ coefficients, meaning that they mice tended to repeat their choice following a failure (Figure 3A, Stay quadrant), while others had negative β_F_ coefficients, indicating a tendency to switch choices following a failure (Figure 3A, Win-Stay, Lose Switch quadrant). The overall observed trial history patterns were similar to that observed in human subjects performing a perceptual decision-making task (Abrahamyan et al. 2016). The second approach evaluated the influence of the history of previous choices on the current choice. The model is equivalent to the model described by (Fründ et al. 2014). This model revealed that the most recent choice had the greatest influence on the current choice (Figure 3B).

**Figure 3.**
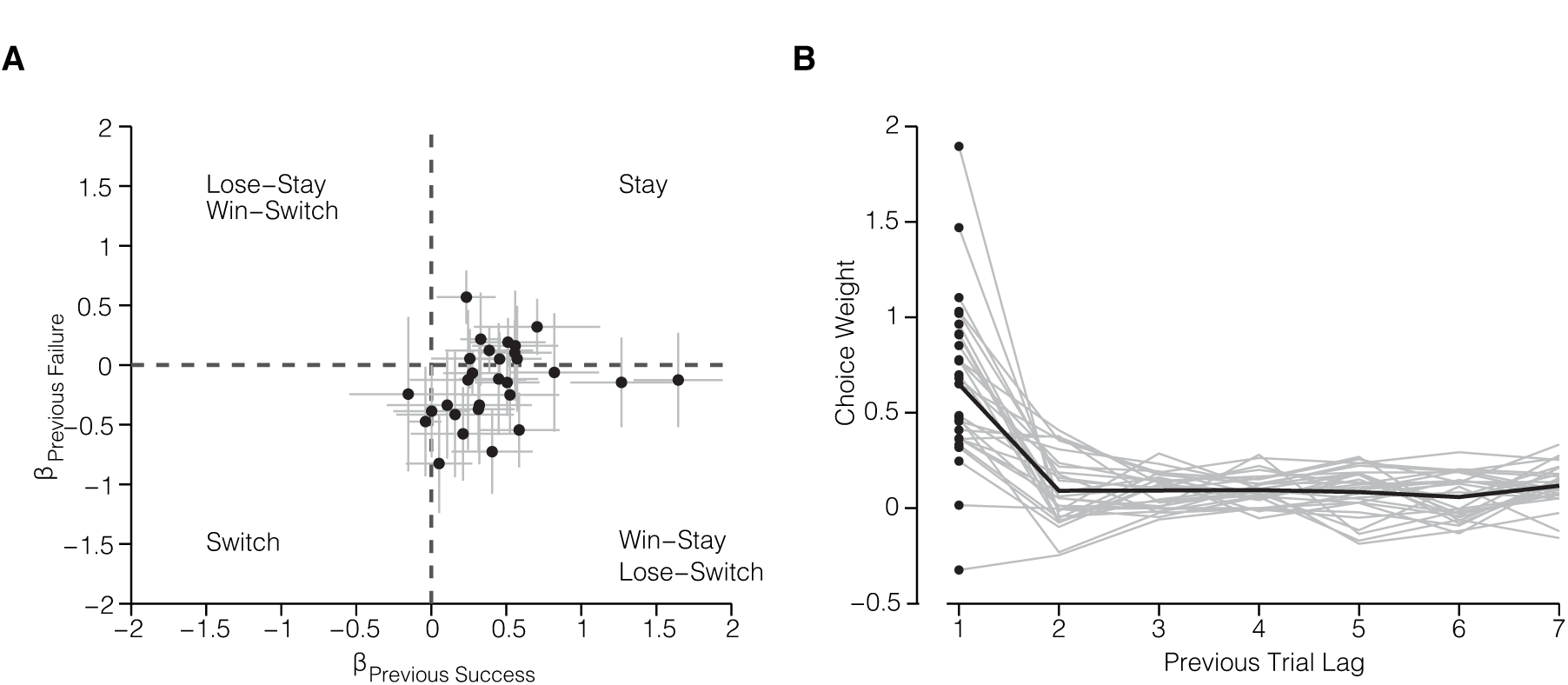
Mice were influenced by both the outcome and choice on the previous trial. (A) Previous choice history: Influence of successes and failures on the current choice for each mouse (*n* = 27 mice). Coefficients were estimated for each session individually and mean coefficients across sessions are plotted. Error bars represent standard error of the mean. (B) Effect of previous choices (*n* = 7 trials in the past). Gray traces, individual subjects (*n* = 27 mice); black trace, average across subjects.

### Decisions are influenced by cumulative brightness

An alternate strategy to accumulating sensory events is to base the decision on the overall brightness experienced over the course of the stimulus. This is a feasible strategy given that the flash event rate is directly proportional to the total LED on-time and therefore the total photons emitted in a sequence.

To test whether mice were influenced by brightness, we performed two brightness manipulation experiments on a subset of animals (Figure 4). First, the intensity of all flashes in a given sequence was randomly increased or decreased on 5% of all trials (Figure 4A). If subjects are influenced by rate alone, their decisions will be the same regardless of brightness (Figure 4B, top row). By contrast if subjects are influenced by brightness, they will report more high-rate choices on brighter trials, and more low-rate choices on dimmer trials (Figure 4B, middle and bottom rows). This is what we observed (Figure 4C, left): brighter stimuli drove a high-rate bias (shift of 1.0±0.5 flashes/s), while dimmer stimuli drove a low-rate bias (shift of 0.8±0.5 flashes/s). When we tested rats on the same manipulation, the changes were even larger (Figure 4C, right): we observed a high rate bias of 4.8±0.9 flashes/s for brighter stimuli and a low rate bias of 3.1 ±0.8 flashes/s for dimmer stimuli.

**Figure 4.**
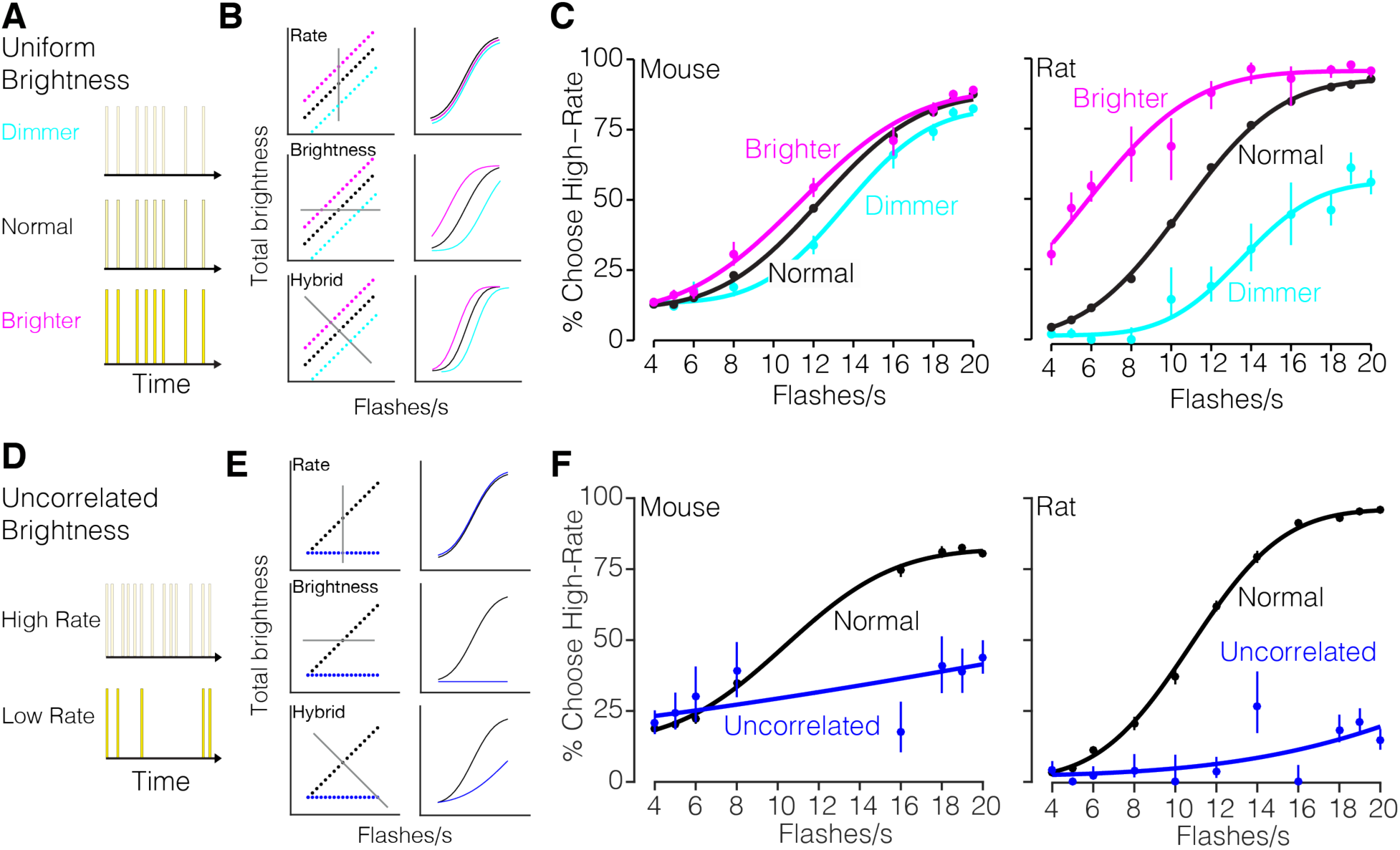
Stimulus brightness influences rate decisions. (A) Schematic of the uniform brightness manipulation experiment. The intensity of individual flashes was varied such that all flashes were dimmer or brighter than normal on 5% of randomly selected trials. (B) Left: stimulus spaces and decision planes (gray lines). Right: Predicted psychometric functions. Each row reflects a candidate way in which the stimulus in each condition would influence decisions given the strategy indicated in the label. Top: stimulus rate. Middle: stimulus brightness. Bottom: Hybrid strategy in which both features are used. (C) Measured psychometric functions. Left: 8 mice; 108,547 trials. Right: 2 rats; 26,201 trials. Points: subjects’ responses. Solid line: 4-parameter cumulative Normal psychometric function fit to the data. Error bars: Wilson binomial 95% confidence intervals. (D) Schematic of uncorrelated brightness manipulation experiment. The intensity of individual flashes was scaled inversely with the flash rate on 5% of randomly selected trials. All sequences have the same cumulative brightness, independent of flash rate. (E) Same as B but for the manipulation in D. (F) Measured psychometric functions. Left: 2 mice; 6326 trials. Right: 2 rats; 9946 trials. Points: subjects’ responses. Solid line: 4-parameter cumulative Normal psychometric function fit to the data. Error bars: Wilson binomial 95% confidence intervals.

Second, we removed the correlation between brightness and flash rate by adjusting the flash intensity in each sequence to the flash rate on 5% of trials. As a result, the total brightness over time was the same across all flash rates (Figure 4D). If subjects are influenced by rate alone, their decisions will be unaffected by brightness (Figure 4E, top row). By contrast if subjects are influenced by brightness, they will have a low-rate bias on uncorrelated trials since the brightness level used was that of the lowest rate stimulus (Figure 4E, middle and bottom rows). This is what we observed: both mice and rats had a low-rate bias (shift of 3±8 flashes/s for mice, 21 ±7 for rats, Figure 4F). Importantly, the dependence of decisions on stimulus rate was reduced but still present (Figure 4F, blue lines not completely flat; i.e. Sensitivity>0, p=4×10^−10^ for mice, p=6×10^−15^ for rats, likelihood ratio test, corrected for parameter on boundary). This argues in favor of a hybrid strategy (Figure 4E, bottom row). If animals had used brightness alone, the psychometric functions would have been flat.

### Inactivation of secondary visual area AM biases perceptual decisions

To test whether the evidence accumulation paradigm engaged cortical circuitry, we sought to reversibly silence secondary visual AM in mice during decision-making. To target AM, we performed widefield retinotopic mapping in each mouse (Figure 5). Briefly, we imaged visually evoked activity in awake transgenic mice expressing GCaMP6 in excitatory neurons (Ai93; CamkIIα-tTA; Emx-cre) in response to a vertical or horizontal bar that periodically drifted across the screen in the four cardinal directions (Kalatsky and Stryker 2003; Garret et al. 2014; Zhuang et al. 2017). This procedure enabled generation of phase maps for altitude and azimuth visual space (Figure 5A,B) and subsequently visual field sign maps (Figure 5C), which were used to estimate the borders between cortical visual areas and reliably identify AM (Figure 5D). Note that the location of area AM is very close to the stereotaxic coordinates used to target PPC (Harvey et al. 2012, Hwang et al. 2017, Goard et al. 2016). Future studies are needed to establish whether AM is a separate cortical area from PPC or whether there is overlap (partial or complete) between the two.

**Figure 5.**
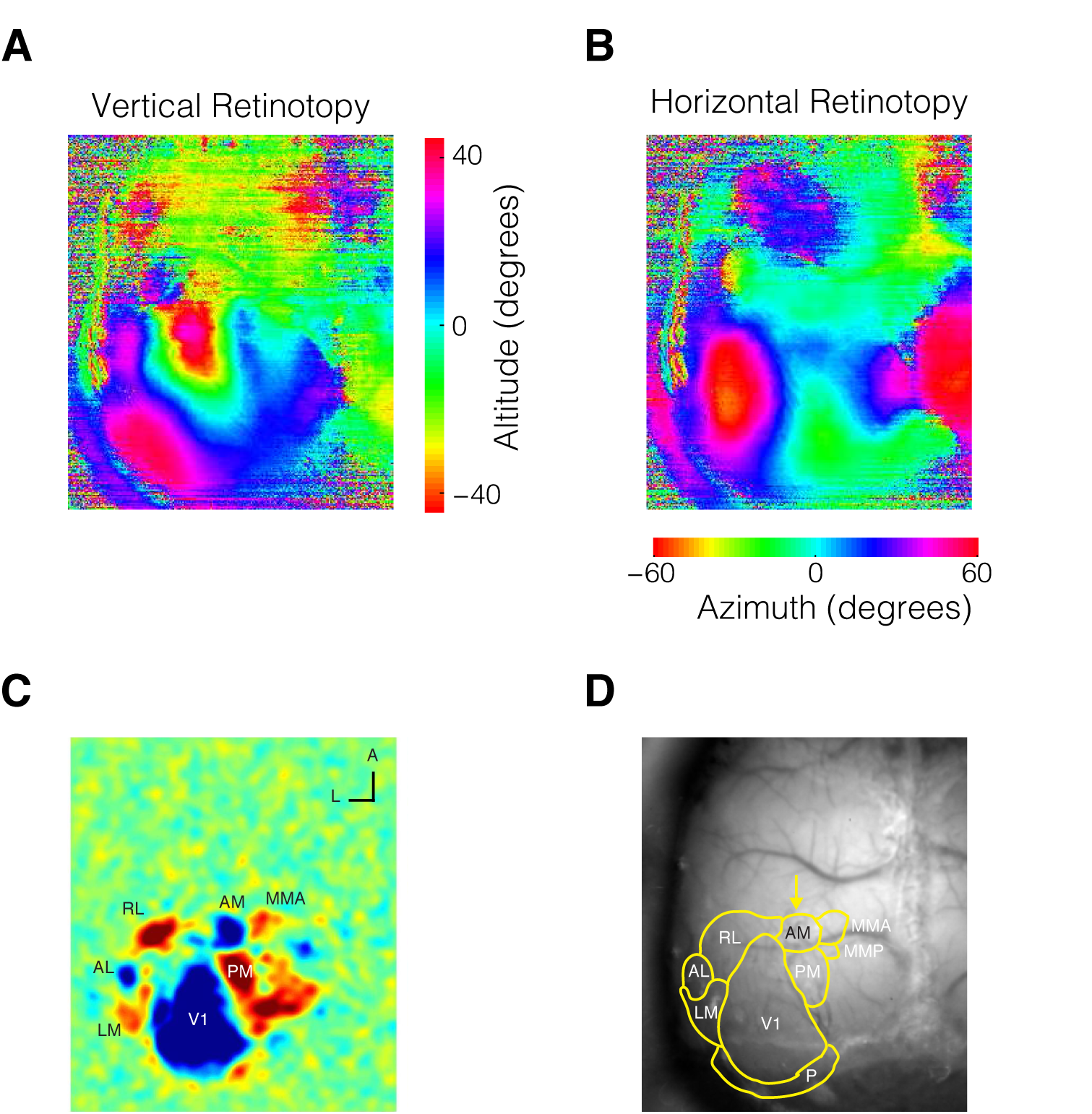
Retinotopic Mapping allows precise localization of visual areas for subsequent manipulation. (A) Altitude and (B) azimuth phase maps (C) Visual field sign map with labeled visual areas (D) Visual area borders overlaid on photograph of skull.

To reversibly silence AM, we used cruxhalorhodopsin JAWS (Halo57), a red light-driven chloride ion pump capable of powerful optogenetic inhibition (Chuong et al. 2014; Acker et al. 2016). Optogenetic stimulation was randomly interleaved on 25% of trials within a session. The optogenetic stimulus pattern consisted of a 1 s long square wave followed by a 0.25 s long linear downward ramp to reduce the effect of rebound excitation that may occur after strong inhibition (Figure 6A, red line) (Chuong et al. 2014; Guo et al. 2014a).

**Figure 6.**
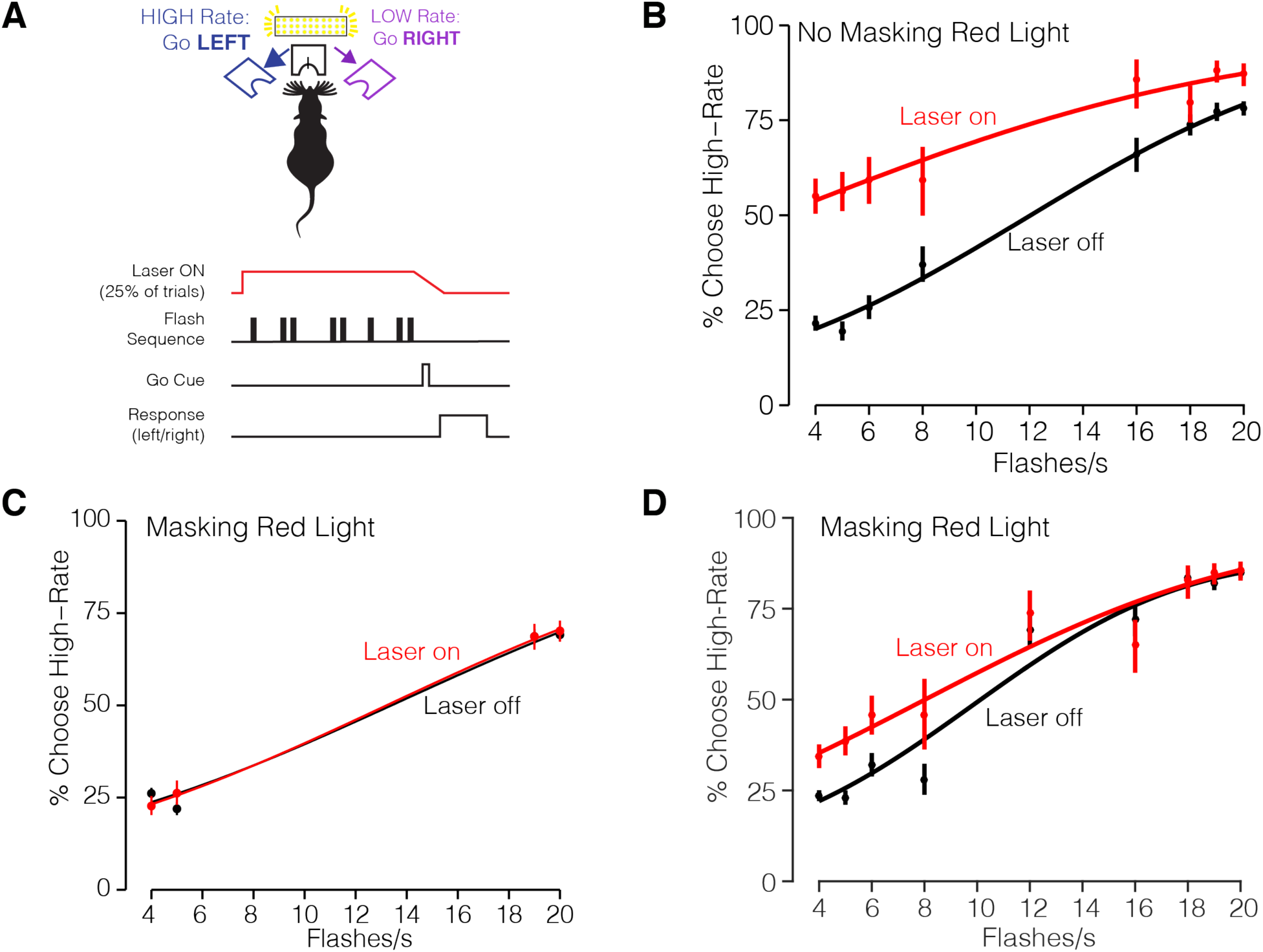
Long wavelength laser stimulation biases decisions in control animals. A) Experimental configuration. Mice were injected with AAV-GFP and implanted with a fiber in right hemisphere area AM. (B) Psychometric function without masking red light (2 mice, 2011 Laser-off trials; 610 Laser-on trials). Irradiance was 32 mW/mm^2^. (C) Psychometric performance with masking red light (2 mice) with easiest flash rate conditions (2699 Laser-off trials; 823 Laser-on trials) (D) Same as C but for sessions including multiple flash rates (2866 Laser-off trials; 903 Laser-on trials). Irradiance was 64 mW/mm^2^.

A potential confound when using JAWS for optogenetic inhibition is that the presence of red light alone may influence behavior. While it was long assumed that rodents are unable to perceive red light, a recent study showed that red-light delivery in the brain can activate the retina and influence behavior (Danskin et al. 2015). Having demonstrated here that decisions can be influenced by brightness (Figure 4), we feared that red-light stimulation might induce a perceived increase in brightness and thus a behavioral bias towards high-rate decisions on photoinhibition trials. To test whether decisions were affected by the presence of red light in the absence of JAWS, we implanted and trained mice injected with a sham virus (AAV-GFP) in AM.

In vivo red light stimulation of sham-injected mice resulted in a high-rate bias (Figure 6B). This confirms the hypothesis that the red light alone influences decisions. The bias is likely because the red lights increased the perceived brightness of the stimulus, driving the animal to make more high-rate choices. Importantly, choices were biased away from the site of the implant, arguing against phosphenes that drew the animal’s attention towards the stimulation side. To counter the red-light bias, we installed additional red lights in the behavior booth to adapt long-wavelength sensitive photoreceptors (Danskin et al. 2015). These external “house lights” strongly reduced the effect of the laser stimulation on behavior (Figure 6C,D).

Although the “house lights” reduced the red-light bias in uninjected animals, the presence of a residual red-light bias in any individual injected animal is difficult to ascertain. If the house-lights were incompletely effective in masking the red-light, the bias could diminish or possibly enhance the effects of direct neural manipulation, depending on how the red-light bias and neural manipulation interact. To surmount this problem, we developed an experimental design in which the stimulus-response contingency was varied while the stimulated hemisphere was held constant. Specifically, we trained two groups of mice on opposing behavioral contingencies: Group A was trained on the contingency: **High-Rate**, go **LEFT**; Group B was trained on the reverse: **High-Rate**, go **RIGHT** (Figure 7A). Both groups were implanted on the left.

**Figure 7.**
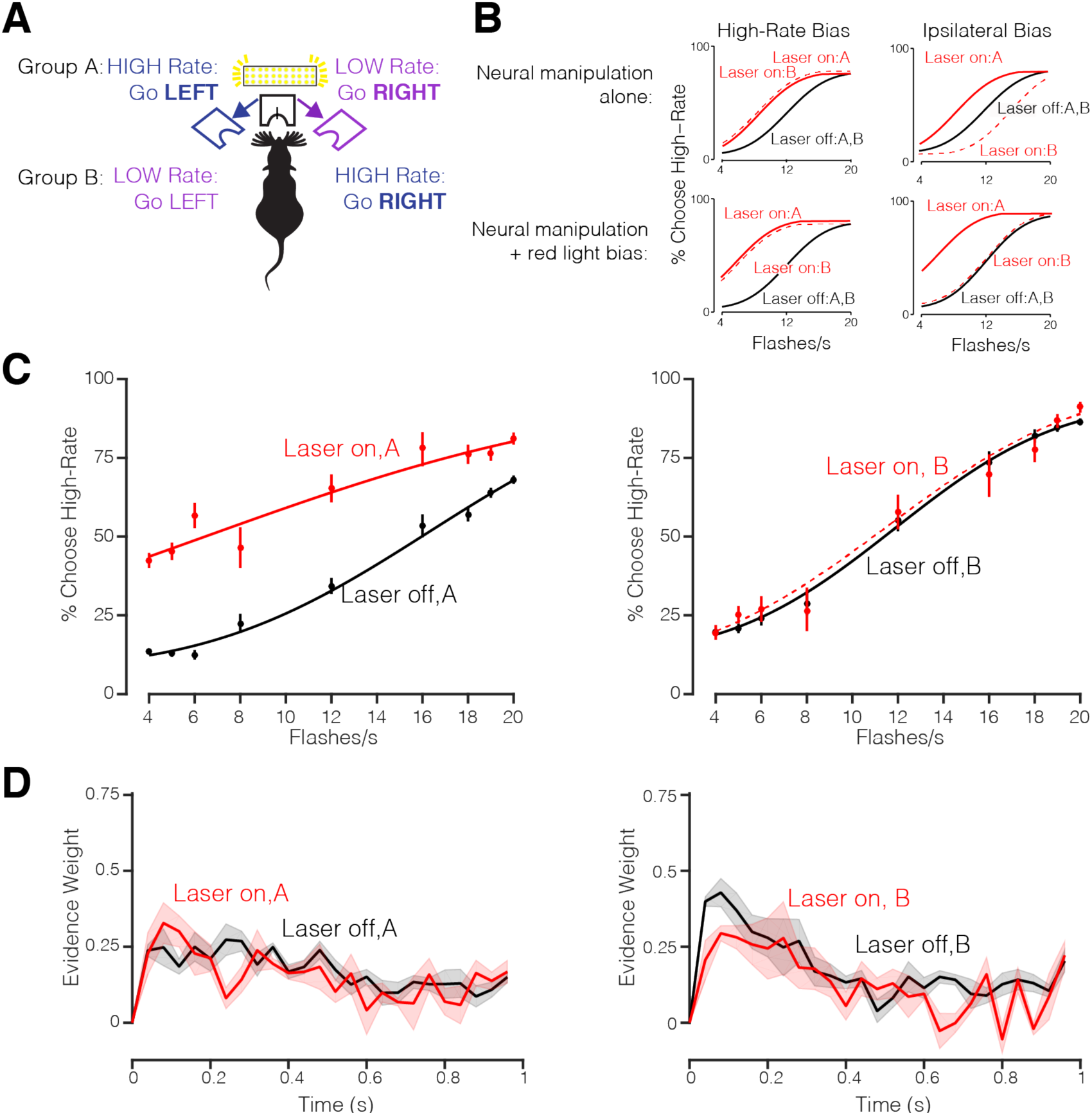
JAWS Photoinhibition of Visual Area AM. (A) Schematic of experimental configuration of AM photoinhibition experiment. Group A mice trained on the contingency: High-Rate, go LEFT and Group B mice trained on the reverse contingency: High-Rate, go RIGHT. Both groups of mice were injected with JAWS virus and implanted with an optical fiber on the left hemisphere. Photoinhibition occurred on 25% of trials during the stimulus period. (B) Predicted behavioral outcomes of AM photoinhibition. Top: predictions for neural manipulation alone. Left: If AM inhibition drives a high-rate bias, the outcomes would be similar for the two groups. Right: If AM inhibition drives a bias towards the ipsilateral side, groups A and B would show biases in opposite directions since the high-rate side differs for groups A and B. Bottom: Predictions for neural manipulation alongside a bias driven by the presence of visible light (as in Figure 6). Left: If AM inhibition drives a high-rate bias, both groups would again exhibit the same bias. Right: If AM inhibition drives an ipsilateral bias, groups A and B would again show biases in opposite directions; potentially with a very weak effect for Group B since the red light and neural manipulations are in opposition. (C,) Decisions for Group A (left, 4 mice; n=5722 Laser OFF trials and n=1958 Laser ON trials) and Group B (right, 3 mice; n=4404 Laser OFF trials and n=1381 Laser ON trials) at laser power irradiance of 64 mW/mm^2^. Circles represent the subject's behavioral response during laser OFF (black) and laser ON (red) trials. Solid line represents the psychometric function fit to cumulative Normal. Error bars represent Wilson binomial (95%) confidence intervals. (D) Left: Psychophysical kernels for group A (37,025 Laser off trials; 12,952 Laser on trials). Right: Psychophysical kernels for group B (32,936 Laser off trials; 12,423 Laser on trials).

One scenario would be difficult to interpret: specifically, JAWS suppression could bias the animal’s estimate of a sensory parameter: rate or perceived brightness for example. If so, the (e.g., high) rate bias from the neural manipulation and the high rate bias from the red-light would be combined similarly in both groups of animals, simply changing the magnitude of the effect in both groups (Figure 7B, left; solid and dashed lines are similar in both top and bottom panels). Distinguishing the effect of the neural manipulation from that of the red-light would be difficult.

Fortunately, in an alternate scenario, JAWS suppression could drive a bias to the ipsilateral side that is independent of the rate associated with that side. In this scenario, the effects will differ for the two groups because the ipsilateral side is associated with high-rate for Group A and low-rate for group B (Figure 7B, top right). The difference across groups will be present even if a red-light bias persists (solid and dashed lines differ in both top and bottom panels in 7B, right). This because the red-light bias will increase high rate choices and thus simply shift both curves leftwards, leaving the difference between them unchanged. For Group B animals, a contralateral (high rate) bias from the red light, combined with an ipsilateral (low rate) bias from the JAWS suppression may potentially cancel each other out (Figure 7B, lower right), leading to an interesting scenario in which the effect of JAWS suppression is only apparent in one group. A comparison of the two groups is therefore essential in interpreting the dual effects of the red-light bias and the JAWS suppression.

A comparison of effects in Group A and Group B revealed a striking difference. Specifically, group A (**High-Rate**, go **LEFT**) had a large, high rate bias (Figure 7C, choice bias = 10.09 flashes/s [2.84 18.72]) accompanied by a change in the slope of the psychometric functions (*β_evidence.opto_* = - 0.045 [-0.064 -0.026] p = 3.4e-06, GLMM Test). However, group B (**High-Rate**, go **RIGHT**) showed no effect (Figure 7D, *β_evidence.opto_* = 0.008 [-0.006 0.022], p= 0.27; choice bias = 0.43 flashes/s [-0.75 1.62]). The group difference was present despite matched laser power, injection volume, and percentage of disruption trials. These data most resemble the lower right panel in Figure 7B, in which the two effects drive the psychometric function in opposing directions: the red light drives a high rate bias, and the JAWS suppression drives an ipsilateral bias, largely canceling each other out. In keeping with the hypothesis that AM suppression drives a change in bias, rather than a change in sensitivity, the psychophysical kernals were largely unchanged in either group of animals (Figure 7D).

## Discussion

We describe a quantitative behavioral paradigm for studying visual evidence accumulation decisions in freely behaving mice. Mice trained on our paradigm performed hundreds of trials per session and maintained stable performance across sessions. A dataset of over half a million trials allowed us to characterize the timecourse of accumulation with precision. We observed that mice were influenced by visual evidence presented throughout the trial, but that they assigned more weight on average to flashes presented earlier in the sequence, similar to monkeys, but unlike rats. Further, despite overall high accuracy, decisions were nonetheless influenced by additional information, such as stimulus brightness and previous reward and choice history. In addition, we demonstrated that area AM plays a causal role in visual decisions. Our experimental design was key in allowing this conclusion because control experiments demonstrated that the red stimulation light biases mice even in the absence of JAWS. Taken together, these results fill a gap in our understanding of accumulation of evidence behavior in mice, and begin to define the causal circuit that supports this behavior.

Our observations of overweighting of information early in the trial are consistent with results from the evidence accumulation paradigm from Morcos and Harvey (2016) in head-fixed mice. Our large cohort of animals and stochastic stimulus arrival times allowed us to more fully characterize this function and extend it to a new paradigm. Interestingly, the shapes of the psychophysical kernels we observed in the mice are qualitatively similar to those observed in nonhuman primates (Katz et al. 2016; Yates et al. 2017). The shape of the kernel differed from that observed in rats trained on the same task (Figure 2B) and previously reported by other evidence accumulation paradigms (Raposo et al. 2012; Brunton et al. 2013; Sheppard et al. 2013; Scott et al. 2015). The difference in psychophysical weighing of evidence across species is intriguing because it suggests that although different species achieve comparable levels of performance, their internal behavioral strategies may differ. This underscores the importance of using stochastic stimuli, which make it possible to uncover the animal’s strategy (Churchland and Kiani 2016).

Our brightness manipulations revealed that decisions were not based solely on rate. For both mice and rats, the cumulative brightness of the flash sequence also influenced decisions (Figure 4C,F). Incorporating brightness in decision-making reflects a clever strategy because in almost all trials, brightness and rate provide evidence in favor of the same decision. In fact, combining these two sources of information is the optimal strategy (Figure 4B,E, bottom row), in the same way that combining auditory and visual information is the optimal strategy in multisensory experiments (Raposo et al., 2012, Sheppard et al., 2013). It would be surprising if animals elected to marginalize brightness when it is such a useful source of information.

The influence of brightness that we observed here contrasts findings from a recent study, which reported that rats performing a visual evidence accumulation task counted individual flashes rather than cumulative flash on-time (Scott et al., 2015). Two experimental design features may explain the difference in results. First, the inter-flash interval in Scott et al. was more than an order of magnitude longer than that used here (250 ms vs. 20 ms) which may discourage a cumulative brightness strategy. Second, Scott et al. didn’t manipulate brightness on catch trials, as we did here, but instead conducted separate experiments in which all trials contained jittered light on-times. This raises the interesting possibility that in Scott et al., the jittered on-times increased the uncertainty of cumulative on-time information. The increased uncertainty would change the optimal strategy, leading animals to weight cumulative on-time less than they would in other experiments in which brightness and event number are correlated.

To understand the neural mechanisms that enable perceptual decision-making, we tested the causal role of secondary visual area AM. Our results, an ipsilateral bias on inactivation trials, suggest that AM normally drives contralateral choices. However, additional studies are needed to more definitively establish the role of AM. Because we observed that red light alone biases choices (Figure 6B), we inferred that the red light and the optogenetic suppression may, in some configurations, bias decisions in opposite directions and cancel each other out (Figure 7C, right). The use of other wavelengths of light to suppress activity (Lien and Scanziani 2013) or the use of non-optical suppression methods such as muscimol (Raposo et al. 2014; Erlich et al. 2015) could provide additional evidence about the role of area AM in decision-making.

Regardless, our results make clear the need to carefully control for light-induced artifacts, both by adapting the animal and by experimental design that disentangles light-induced artifacts from true behavioral changes. The artifact we observed is most likely caused by red light propagating from the stimulation site through the brain and directly activating the retina. Danskin et al. (2015) measured retinal activation during in vivo red light stimulation and found the largest activation ipsilateral to the implanted stimulation fiber. Interestingly, the bias that we observed here was contralateral to the implanted stimulation fiber. Notably, in this cohort, the contralateral side was associated with high-rate choices (Fig. 6A). A likely explanation is that light from the fiber increased overall brightness; as we demonstrated in separate experiments (Figure 4), increased brightness can be interpreted as evidence for high rate choices. Additional experiments that systematically vary the stimulus response contingency in sham-injected animals could confirm this hypothesis.

The proposed ipsilateral bias caused by AM photoinhibition is consistent with spatial hemineglect observed in visual parietal lesions. Spatial hemineglect, also referred to as contralateral neglect, is a phenomenon that occurs when subjects ignore the contralateral hemifield as a result of lesion to the parietal cortex. Although hemineglect has been reported in humans (Stone et al. 1991; Kerkho 2001) and rats (Crowne et al. 1986; Reep and Corwin 2009), we could not find a report on mice. The presence of hemispatial neglect would suggest that the mice are neglecting the tendency to go towards the affected (contralateral) visual hemifield. A related interpretation of the ipsilateral bias due to suppression of AM activity is that neurons in AM are active in advance of contralateral choices. In this scenario, the two hemispheres of AM would represent competing movement intentions, such that inactivation of one hemisphere leads to movement in the opposing direction.

Taken together, these observations begin to address two major gaps in our understanding of accumulation of evidence behavior: we have precisely characterized the timecourse of evidence accumulation, and have uncovered that rate, brightness, and trial history jointly shape decisions. Finally, our results support a role for AM in visually guided evidence accumulation decisions. We propose that AM drives contralateral choices in the visual flashes task, such that AM inhibition leads to an ipsilateral bias. This is consistent with anatomical projections of AM to motor areas (Wang et al. 2012; Allen Brain Atlas 2015) and the recently proposed role for mouse parietal cortex in navigation (Krumin et al. 2017). These results point to the need for more causal manipulations in mouse visual areas, and highlight the challenge in designing experiments and interpreting data when light is used to transiently manipulate neural activity.

## Acknowledgements

We thank Rob Eifert and Barry Burbach for technical support. We thank Simon Musall, Matthew Kaufman, Ashley Juavinett, and Lindsey Glickfeld for helpful discussions and feedback on the manuscript. This work was supported by NIH NRSA F31 (1F31-EY025164) to O.O, The Simons Collaboration on the Global Brain, The Klingenstein-Simons Foundation and the Pew Charitable Trust to A.K.C.

